# A hierarchical model for external electrical control of an insect, accounting for inter-individual variation of muscle force properties

**DOI:** 10.1101/2022.12.19.521014

**Authors:** Dai Owaki, Volker Dürr, Josef Schmitz

**Affiliations:** Dept. of Robotics, Graduate School of Engineering, Tohoku University; Dept. of Biological Cybernetics, Faculty of Biology, Bielefeld University; Centre for Cognitive Interaction Technology, Bielefeld University

## Abstract

Cyborg control of insect movement is promising for developing miniature, high-mobility, and efficient biohybrid robots. However, considering the inter-individual variation of the insect neuromuscular apparatus and its neural control is challenging. We propose a hierarchical model including inter-individual variation of muscle properties of three leg muscles involved in propulsion (retractor coxae), joint stiffness (pro- and retractor coxae), and stance-swing transition (protractor coxae and levator trochanteris) in the stick insect *Carausius morosus*. To estimate mechanical effects induced by external muscle stimulation, the model is based on the systematic evaluation of joint torques as functions of electrical stimulation parameters. A nearly linear relationship between the stimulus burst duration and generated torque was observed. This stimulus-torque characteristic holds for burst durations of up to 500 ms, corresponding to the stance and swing phase durations of medium to fast walking stick insects. Hierarchical Bayesian modeling revealed that linearity of the stimulus-torque characteristic was invariant, with individually varying slopes. Individual prediction of joint torques provides significant benefits for precise cyborg control.

## Introduction

Hybrid insect–computer robots (***Krause et al., 2011***; ***Li and Sato, 2018***) represent cutting-edge approaches to develop robots with locomotor performances comparable to those of insects. With the advancement and diversity in micro-flexible and micro-printable electronics (***Rogers et al., 2010***; ***Rich et al., 2021***), micro-mechanical fabrication, and micro-actuator technologies (***Kim et al., 2020***), such biohybrid, i.e. cyborg robots have been engineered to manipulate their gait and flight through electrical stimulation of target muscles in various insects, includings beetles (***Sato et al., 2009***; ***Sato and Maharbiz, 2010***; ***Sato et al., 2015***; ***Cao et al., 2016***; ***Doan et al., 2018***; ***Nguyen et al., 2020***; ***Kosaka et al., 2021***), moths (***Sane et al., 2007***; ***Bozkurt et al., 2009***; ***Hinterwirth et al., 2012***; ***Ando and Kanzaki, 2017***), and cockroaches (***Sanchez et al., 2015***). The advantage of biohybrid (cyborg) robots is that they do not require individual “design,” “fabrication,” and “assembly” processes for each component because they use the body tissues of living insects (***Cao et al., 2014***). Moreover, cyborg robots have low power consumption, that is, a few milliwatts (***Sato and Maharbiz, 2010***). Although studies on insect cyborgs have demonstrated simple manufacturing and promising energy efficiency, they are still in the initial phase of development from the perspective of evaluating both feasibility and reliability of their control.

Perhaps the greatest challenge in cyborg control comes with the inter-individual variability of animals. Past neurophysiological studies related to animal neural activity have discussed the failure of averaging-based approaches, in which a model formulated using the average data for a group cannot explain the characteristics of any individual in the group (***Golowasch et al., 2002***; ***Schulz et al., 2006***). For example, variable and non-periodic patterns in feeding behavior of Aplysia have been reported to be subject to strong inter-individual variation (***Horn et al., 2004***; ***Brezina et al., 2005***; ***Zhurov et al., 2005***). In insect motor physiology, the prediction error of muscle models which are based on sample averages is very high (***Blümel et al., 2012c***) and may be halved using individual-specific model (***Blümel et al., 2012a***). At the level of leg movements, variability has been investigated in lobsters (***Thuma et al., 2003***) and stick insects (***Hooper et al., 2006***). The variability of whole-body locomotion arises from step parameter variation of single legs (***Theunissen and Dürr, 2013***) but also from variation of coupling strength among legs (***Dürr, 2005***). One possible approach for accounting for inter-individual variability in cyborg control of single-leg movement is to construct a feedback control system (***Cao et al., 2014***). Although the kinematics-control of joint anges (***Cao et al., 2014***) has exhibited remarkable performance, its applicability to the control of dynamic gaits, such as that for walking, is still controversial. Furthermore, insects have abundant control variables, that is, degrees of freedom in their actuators and sensors. At present, the number of control variables of current insect cyborgs has to be reduced2 owing to system implementation difficulties.

A promising approach to overcome the “pitfalls” associated with averaging across individuals is to understand the underlying principles that govern inter-individual variability in insect motor control. Especially, the output characteristics of muscle are key for controlling the dynamics of movement: muscles convert neural activity into movement and then generate behavior from interactions with the environment. In conjunction with current models of muscle activation (***Harischandra et al., 2019***) and contraction dynamics (***Blümel et al., 2012b***), we can exploit experimental data to tell parameters that are strongly influenced by inter-individual variation as opposed to others that are common characteristics. To this end, we employed a hierarchical modeling framework based on the Bayesian statistical analysis (***Watanabe, 2018***; ***Gelman et al., 2013***) that explicitly accounts for inter-individual variation in experimental data. In particular, we applied a set of hierarchical Bayesian models with different combinations of common and individually varying parameters and mathematically evaluated their prediction performance.

The main objective of this study was to systematically evaluate how muscle force and corre-sponding joint torques depend on external electrical stimulation, as a fundamental pre-requisite for precise insect cyborg control. To this end, we measured joint torques induced by stimulating one out of three leg muscles in the middle leg of the stick insect species *Carausius morosus* (de Sinéty, 1901): these were the protractor coxae, retractor coxae, and levator trochanteris. We focused on these three proximal muscles because the retractor coxae is the primary muscle for propulsion34, the pro/retractor coxae contributes to weight-dependent postural adjustment by regulating joint stiffness (***Dallmann et al., 2019***; ***Günzel et al., 2022***), and the levator trochanteris is important for postural termination and swing initiation (***Dallmann et al., 2017***). Using a custom-built electrical stimulator to generate parameter-tunable pulse-width-modulated (PWM) signals, we simulated burst-like activity of motor neurons in insects and measured the corresponding joint torques generated in response to our electrical stimuli. Using Bayesian statistical modeling and the “widely applied information criterion” (WAIC) index (***Watanabe, 2018***) for model prediction, we evaluated several model variants to identify the one that explained the experimental data best. In particular, we evaluated the predictive performances of model variants with and without interindividual variation of experimental parameter estimates. A piecewise linear relationship was observed between the burst duration and the joint torque generated for a given parameter set of the PWM burst. Linearity was found to hold for burst durations of up to 500 ms, which corresponds to the stance phase (300 to 500 ms) and swing phase (to 250 ms) of a stick insect walking at medium to fast speeds (***Dürr et al., 2018***). Furthermore, the hierarchical Bayesian modeling revealed both invariant and individually varying characteristics of joint torque generation in stick insects. This allows for individual tuning of electrical stimulation parameters for highly precise insect cyborg control.

## Results

### Burst duration and generated joint torque

Figure 1 illustrates the obtained relation between the PWM burst duration and the generated joint torques for the protractor (A), retractor (B), and levator (C) muscles from 10 animals (*N* = 10). The parameters of the PWM signals were set to 2.0 V, 50 Hz, and 30% duty ratio. During one trial, we stimulated one muscle n times with fixed PWM parameters and measured the generated torque at the corresponding joint.

**Figure 1.**
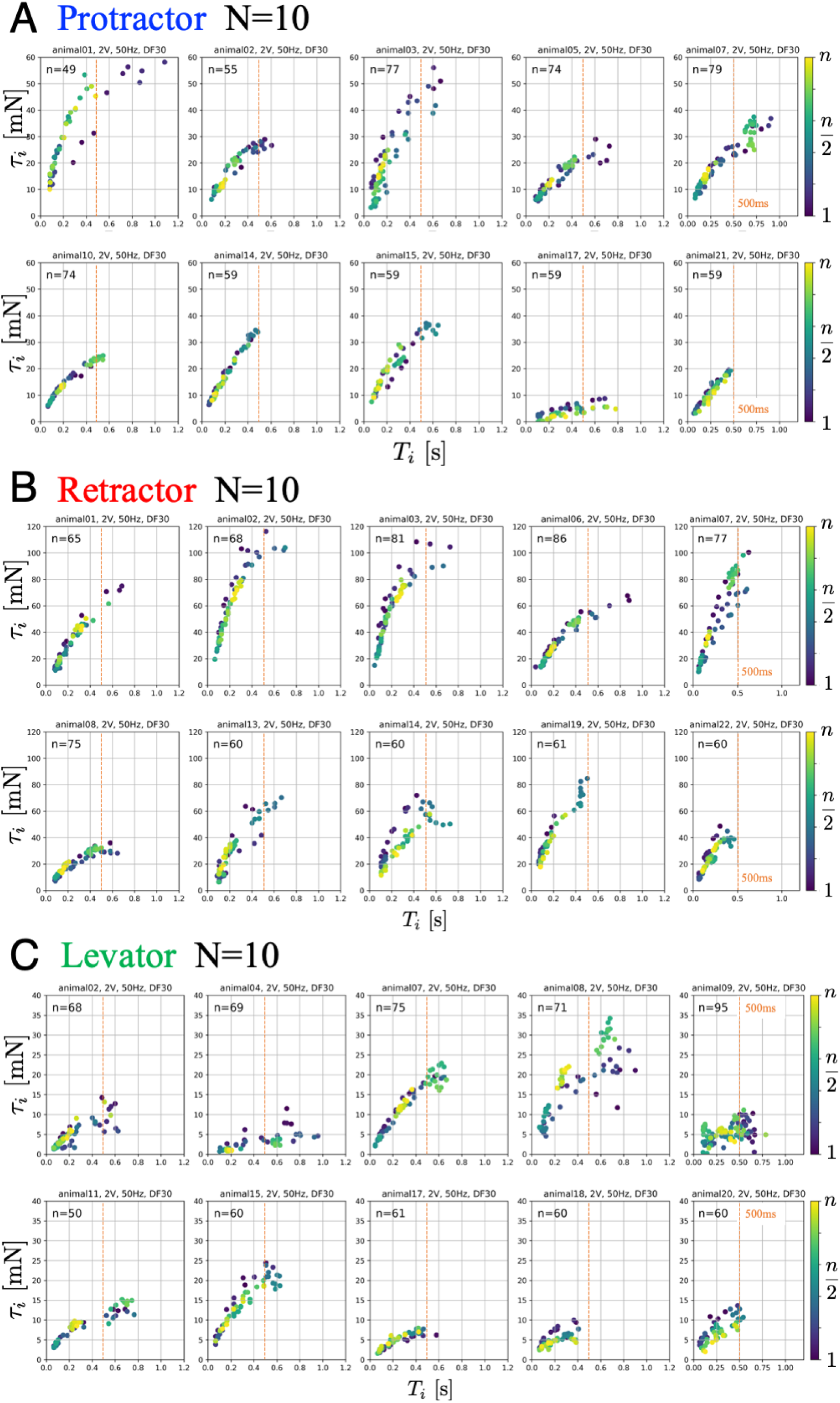
Joint torques as a function of burst duration. Data from 10 animals with three muscles, each: (**A**) protractor, (**B**) retractor, and (**C**) levator muscle stimulation. The PWM burst parameters were 2.0 V. 50 Hz, and 30% duty ratio. *n* gives the number of stimulations for each animal. The color of the symbols indicates the order of the stimulations: blue (1) to yellow (*n*). The positive values of joint torque represent intended (**A**) forward, (**B**) backward and (**C**) upward rotation of the coxa relative to the thorax.

The results indicate the input–output relation (burst duration and generated torque) corresponded to a linear function or a power function with an exponent of less than 1.0. Furthermore, the relationship holds for burst durations up to 500 ms for all animals, corresponding to the duration of swing and stance phases in medium to fast walking stick insects (***Dürr et al., 2018***). Maximum torques for ThC and CTr joints were 60 mN-m, 120 mN-m, and 40 mN-m (***Dallmann et al., 2019***). For a given set pf PWM parameters, the generated torques remained almost constant for all stimulations, suggesting that muscle fatigue to be negligible for at least *n* = 50 stimulations.

### Comparison of model predictability

Using the WAIC described in the previous section, we compared the prediction performance of the six models. Figure 2 (A)-(C) shows the WAIC values for each voltage applied (1.0–4.0 V) and models 1-1 to 2-4. Fig. 2 (A)-(C) show the results for the protractor, retractor, and levator muscles, respectively. For stimulation experiments on each of the three muscles, the models with a hierarchical parameter for expressing individual differences for 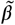 (models 1-2 and 2-2) had the lowest WAIC and, therefore, the best predictive performance. Conversely, the model with individual differences for both 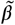 and 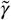 (model 2-4) exhibited the lowest prediction performance, indicating that inter-individual variation of the exponent does not improve model estimates.

**Figure 2.**
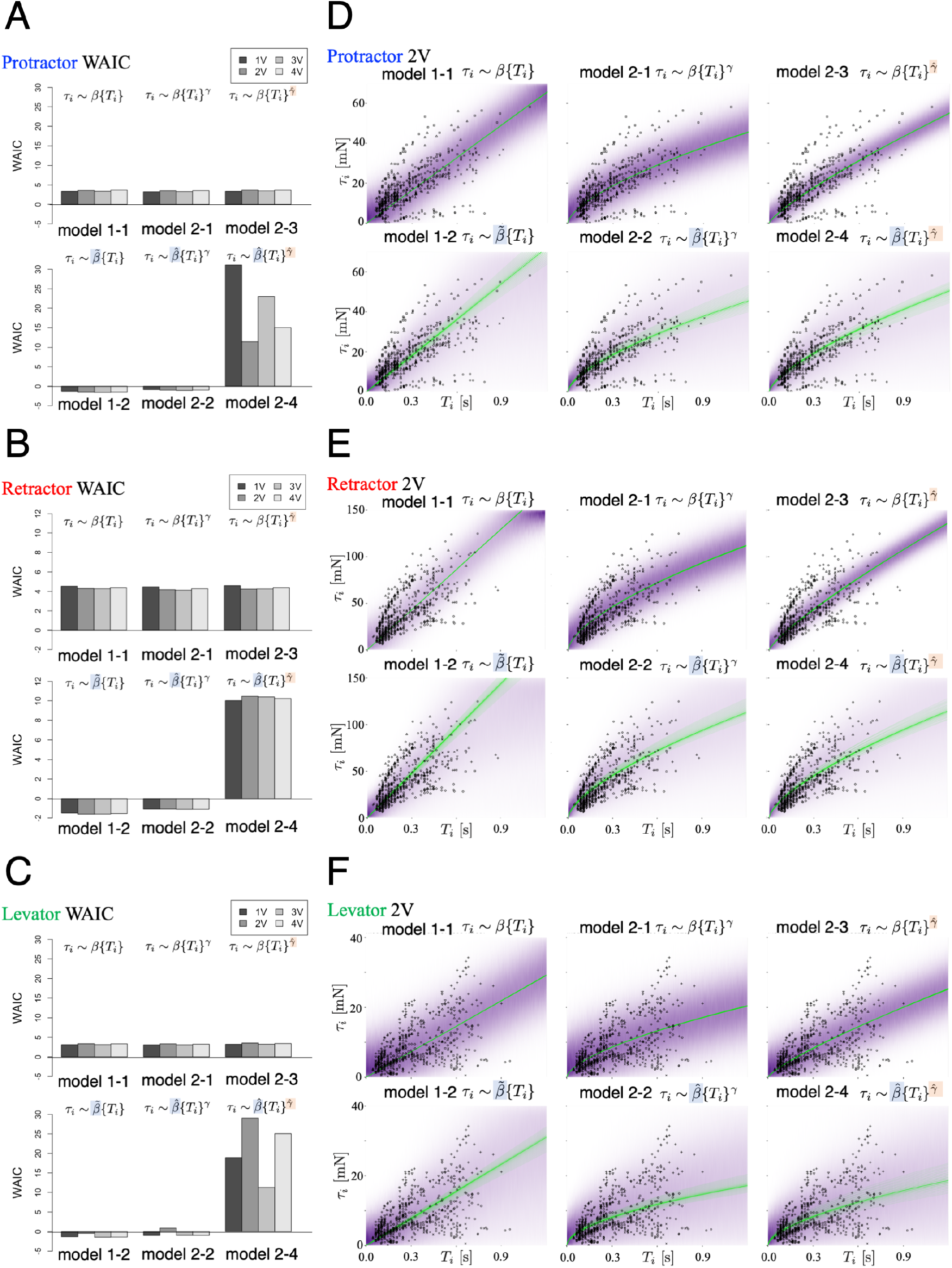
Model comparison underscores significance of inter-individual variation of slope. We compared the six models that were explained in the “Model” subsection. (**A**), (**B**), and (**C**) show plots of the WAIC (***Watanabe, 2018***) values for the protractor, retractor, and levator stimulations, respectively (*V* = 10 animals per muscle). The parameters with tilde, 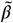 and 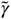, indicate that the parameters include inter-individual variation. PWM parameters were set as follows: (1.0 V, 2.0 V, 3.0 V, and 4.0 V), at 50 Hz and 30% duty ratio. Negative values were obtained for models 1-2 and 2-2 for all voltages and all muscles. The lowest WAIC indicates the best prediction model, as explained in the “WAIC” subsection. Right panels show Bayesian predictive estimation for the protractor (**D**), retractor (**E**), and levator (**F**) stimulation experiments with PWM parameters 2.0 V, 50 Hz, and 30% duty ratio. The differences in the point styles indicate individual animals. In each panel, the violet shading indicates the probability density of the distribution predictive. The green lines represent twenty samples from the posterior distribution in decreasing order of probability density.

### Bayesian estimation of generated torque for a given burst duration

Figure 2 (D)-(F) shows the predictive distributions for data of a new animal using the Bayesian posterior distribution for the six models. The results were obtained with PWM bursts at 2.0 V voltage, 50 Hz frequency, and 30% duty ratio. The results show that the hierarchical models (model 1-2 and model 2-2) for the 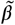 parameter can successfully and adequately capture the range of experimental results on (D) protractor, (E) retractor, and (F) levator torques for all animals. This suggests that, compared with other models, the hierarchical models can appropriately account for interindividual variation of muscle properties for new unknown animals. Figure 3 illustrates this by overlapping the experimental data for 10 animals in Fig 1 with the distributions predicted by the linear hierarchical model (model 1-2) for each individual.

**Figure 3.**
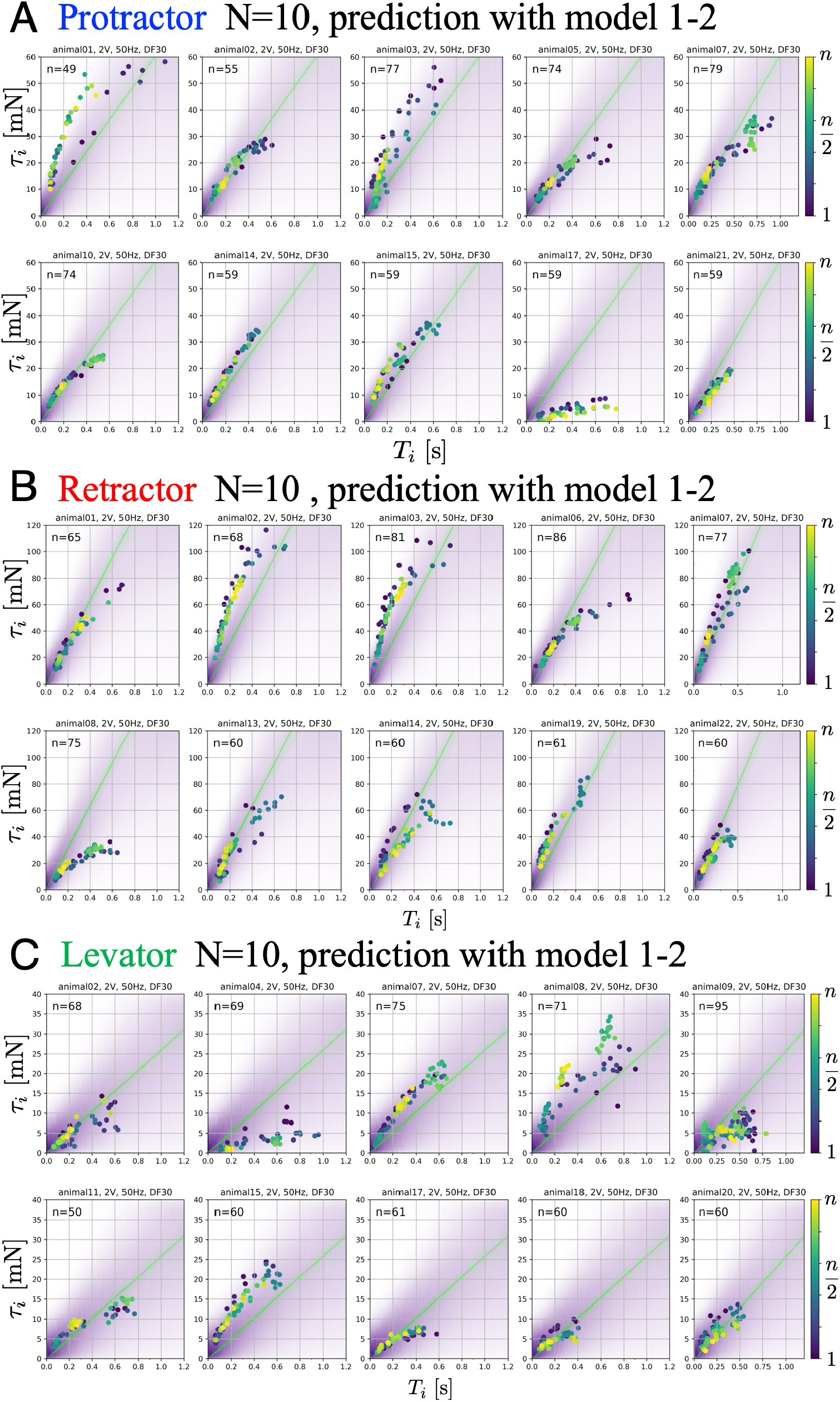
Predictive distributions from the linear hierarchical model (1-2) for each individual: The protractor (**A**), retractor (**B**), and levator (**C**) stimulation experiments with PWM parameters, 2.0 V, 50 Hz, and 30% duty ratio. n gives the number of stimulations for each animal. The color legend indicates the order of the stimulations: blue (1) to yellow (*n*). In each panel, the violet shading indicates the probability density of the predictive distribution. The green lines represent twenty samples from the posterior distribution in decreasing order of probability density. The results demonstrate that the linear hierarchical model had an accurate predictive distribution in the range up to 500 ms.

### Effect of an individual animal and applied voltage on muscle properties

Figure 4 shows the changes in the muscle characteristic parameters *β* and *γ* with respect to changes in the voltage applied. We calculated the changes in *β* and *γ* with respect to the applied voltage from the experimental results, using the six Bayesian models.

**Figure 4.**
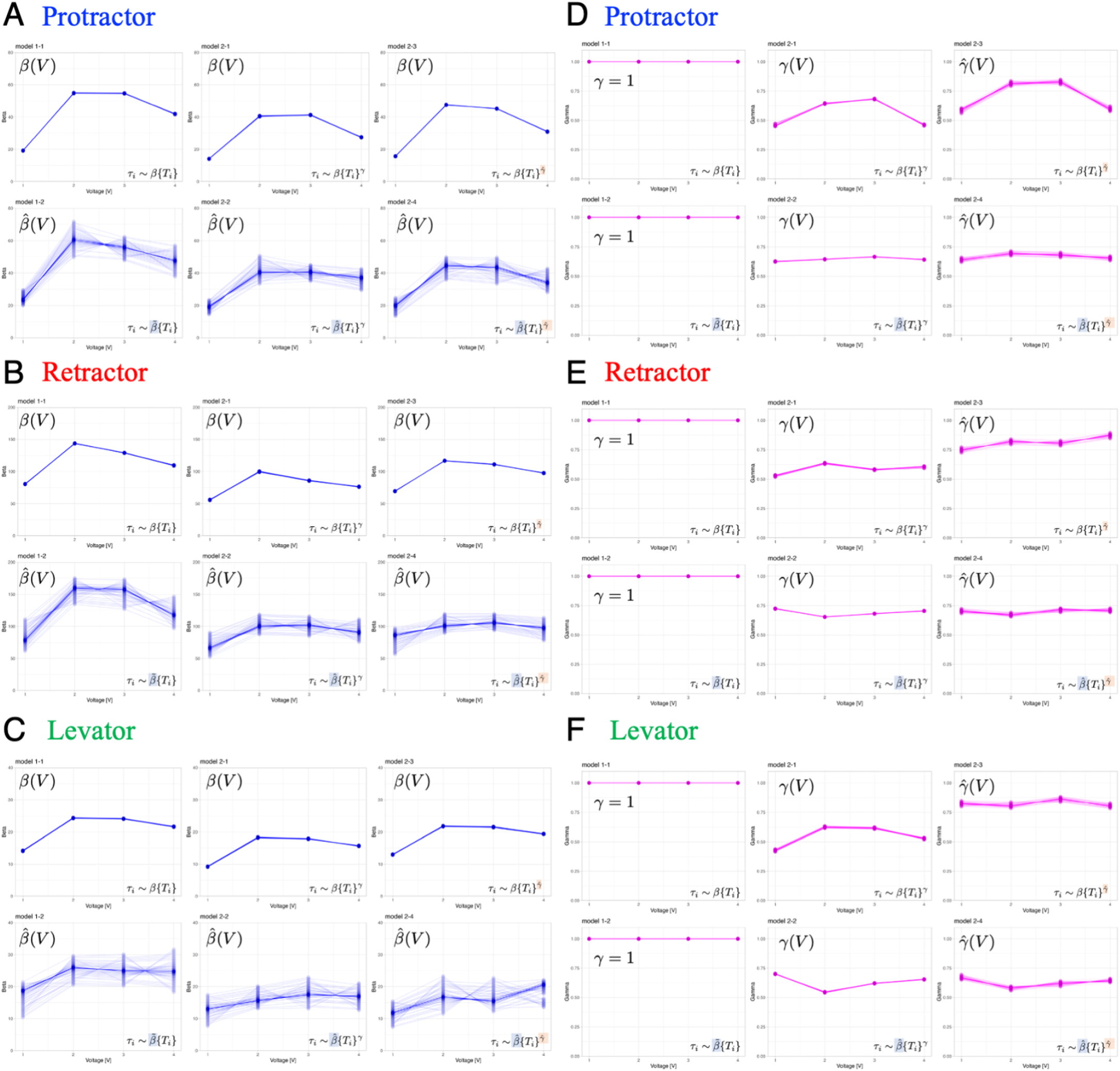
Dependence of muscle property parameters on the applied voltage and individual animals in the six models: The left graphs (**A**), (**B**), and (**C**) represent the estimation for *β* for the applied voltage varied from 1.0 to 4.0 V for the six models. (**A**) and (**D**), (**B**)and (**E**), and (**C**) and (**F**) illustrate the protractor, retractor, and levator stimulations, respectively. In (**A**)to (**C**), the upper and lower panels show non-hierarchical (1-1,2-1, 2-3)and hierarchical (1-2, 2-2, 2-4) models for *β*, respectively. The right graphs (**D**), (**E**), and (**F**) represent the estimation for *γ* in applied voltage changes. In (**D**) to (**F**), the left panel shows linear models (*γ* = 1,1-1,1-2); the middleand right panels illustrate non-hierarchical (2-1, 2-2), and hierarchical (2-3, 2-4) models for *γ*, respectively. For hierarchical models (1-2, 2-2, 2-3, 2-4), the plot includes twenty samples from the posterior distribution in decreasing order of probability density, showing inter-individual variation.

The results indicate the following three points: (1) *β* varied with the applied voltage, and there exists an optimal voltage that maximizes *β*; (2) except for non-hierarchical nonlinear models (models 2-1 and 2-3), *γ* has a low dependence on the applied voltage; and (3) *β* is strongly subject to inter-individual variation (large variability), whereas *γ* is affected much less.

## Discussion

In this study, we investigated externally controlled joint torques induced by external electrical stimulation of one out of three leg muscles (protractor, retractor, and levator) in the stick insect *Carausius morosus*. For a given parameter set for PWM burst stimulation, we found a piecewise linear relationship between the burst duration and generated joint torque. Linearity holds fora burst duration up to 500 ms. Fora more detailed analysis of the joint torques generated by leg muscles, we used Bayesian statistical analysis and modeling to account for inter-individual variation. A comparison of the six models (with combinations of linear, nonlinear, non-hierarchical, and hierarchical models) showed that the two models that include inter-individual variation of slope parameter *β* performed best. Models 1-2 and 2-1 most accurately predicted the posterior predictive distribution.

The exponent *γ* is a macroscopic property of the generated joint torque, that is the degree of non-linearity of the stimulus-torque characteristic; it is linear when *γ* = 1. Conversely, slope parameter *β* defines the rate of increase of the generated torque. In a comparison of the prediction performance of models in Fig. 2, the mathematical index WAIC revealed that the two models in which only *β* was a hierarchical parameter (models 1-2 and 2-1), performed best. Since only hierarchical parameters account for inter-individual variation, we conclude that *β* is strongly affected by individual differences, whereas *γ* is invariant among specimens. Thus, we found that the macroscopic properties of leg muscles are common to all individuals, whereas individuals differ in the slope *β*, i.e. the rate by which the three types of leg muscles respond to electrical stimulation. Furthermore, as shown in Fig. 4, we found that *β* was highly affected by the applied voltage, whereas the exponent *γ* was close to unity, largely independent of the applied voltage, indicating that the macroscopic properties of leg muscles were invariant to the applied voltage. We conclude that linearity was an invariant feature of the stimulus-torque characteristic, whereas the slope of this characteristic varies among individual stick insects and with the applied voltage. These results are in line with those of existing studies on the properties of myogenic forces in other insect species (***Cao et al., 2014***; ***Blümel et al., 2012c***; ***Harischandra et al., 2019***): the generated torque depends much less on PWM voltage and frequency (***Blümel et al., 2012c***; ***Harischandra et al., 2019***) than it depends on burst duration, suggesting the total number of subsequent input pules are important. This is indeed what would be expected for a slow insect muscle (***Blümel et al., 2012c***) that essentially “counts” incoming spikes within a given time window. Compared to the non-linear properties of muscle, we show that our monitoring of torques in an intact animal resulted in a linear characteristic (for intervals up to 500 ms) that would not be expected from isometric force measurements of isolated muscle.

The comparison of the linear model (model 1-2) with the nonlinear model (model 2-2) using the WAIC for all conditions (muscle type and applied voltage) resulted in lower values for the linear model. Models with lower WAIC can generate predictive distributions closer to the true distribution while using fewer parameters (***Watanabe, 2018***), suggesting that the experimental results obtained in this study can be adequately explained using a linear hierarchical Bayesian model (1-2). This model renders it useful for predicting the generated torque for each new animal in real-time during an experiment. Specifically, by assuming the linear hierarchical Bayesian model, we can measure responses to very few PWM stimulus bursts and estimate *β* for the current individual’s stimulus-torque characteristic. This allows an experimenter to acquire an appropriate muscle model of an unknown animal in a short time without having to use potentially time-consuming machine learning methods, such as deep learning algorithms. Moreover, the properties were linear for stimulus burst durations up to 500 ms. This linearity region corresponds to the stance and swing phase durations of medium-speed to fastwalking stick insects of the species *Carausius morosus* (***Dürr et al., 2018***). The magnitudes of the joint torques generated by the protractor, retractor, and levator were comparable to those for resisted movement during stick-insect walking, e.g. coxatrochanterjoint depression during stance (***Dallmann et al., 2016***). This suggests that the estimated stimulus-torque characteristic captures the natural dynamic characteristics of leg muscles during walking, both in terms of duration of excitation as well as maximum torque.

This study takes a first but important step towards highly precise insect cyborg control. In previous studies, we defined Motion Hacking (***Owaki et al., 2019***; ***Owaki and Dürr, 2022***) as a technique for controlling insect leg movements through external electrical stimulation, while retaining the insect’s own nervous system and sensorymotor loops. This approach requires a collaborative effort of engineering and biology in order to elucidate how adaptive walking ability of insects may be exploited for biohybrid control of motor flexibility. The Motion Hacking (***Owaki et al., 2019***; ***Owaki and Dürr, 2022***) method strives to observe the adaptation process in the insect’s own sensorymotor system as leg movements are intentionally controlled by a human operator, so as to reveal hidden mechanisms underlying natural locomotion. Thus far, research on insect cyborg control has addressed aspects of flight control (***Sato et al., 2009***; ***Sato and Maharbiz, 2010***; ***Sato et al., 2015***; ***Kosaka et al., 2021***; ***Sane et al., 2007***; ***Bozkurt et al., 2009***; ***Hinterwirth et al., 2012***), gait control (***Cao et al., 2016***; ***Doan et al., 2018***; ***Nguyen et al., 2020***; ***Ando and Kanzaki, 2017***; ***Sanchez et al., 2015***), and controlling jellyfish propulsion (***Xu and Dabiri, 2020***; ***Xu et al., 2020***). In contrast to our present study, the main objective of the mentioned studies was to convert target animals into cyborgs, with little examination of the control mechanisms and/or muscle properties involved. Here, we used PWM pulse bursts to mimic motor neuron commands during insect locomotion, and selected key muscles to estimate stimulus-torque characteristics reliably and in very short time. Then, we used Bayesian statistical modeling to tell which parameters were subject to interindividual variation and which were not. Our finding of linear characteristics with inter-individual variation of slope show compellingly how a systematic engineering intervention to an otherwise intact animal motor system can yield a simple, technically exploitable description of motor system properties. We argue that this description could not have been obtained by methods addressing isolated neural circuits or partial anatomical structures, but required the physical intactness of the natural system.

Still, there are several limitations to the present study. First, as in many neurophysiological experiments (***Berg et al., 2012***; ***Lepreux et al., 2019***), stick insects were fixed and not walking in the experimental setup (Fig. 5 A). Although there are only few studies on the natural dispersal behavior of stick insects, it is clear that they spend a lot of their lifetime at rest, e.g. in camouflage. Their tendency to attain camouflage postures can be exploited in experiments, as it is relatively easy to restrain active, spontaneous leg movements in an experimental setup. Nevertheless, the possibility to conduct combined motion capture and EMG recordings in freely walking stick insects (***Dallmann et al., 2019***; ***Günzel et al., 2022***; ***Dallmann et al., 2017***; ***Bidaye et al., 2018***) suggests that Motion Hacking during unrestrained, voluntary locomotion will be feasible in the future. Whereas the range of PWM burst duration and the joint torques generated are well within the physiological range, there is still considerable discrepancy between the PWM signals generated by our Raspberry Pi microcontroller and the natural firing patterns of motor neuron pools (***Günzel et al., 2022***). Future research will need to examine how much the simplification of the driving burst input affects the time course of the torque generated. So far, it is re-assuring that the simplified PWM signal used here could be applied more than 50 times in a sequence without causing muscle fatigue, i.e. with a sustained level of generated torque.

**Figure 5.**
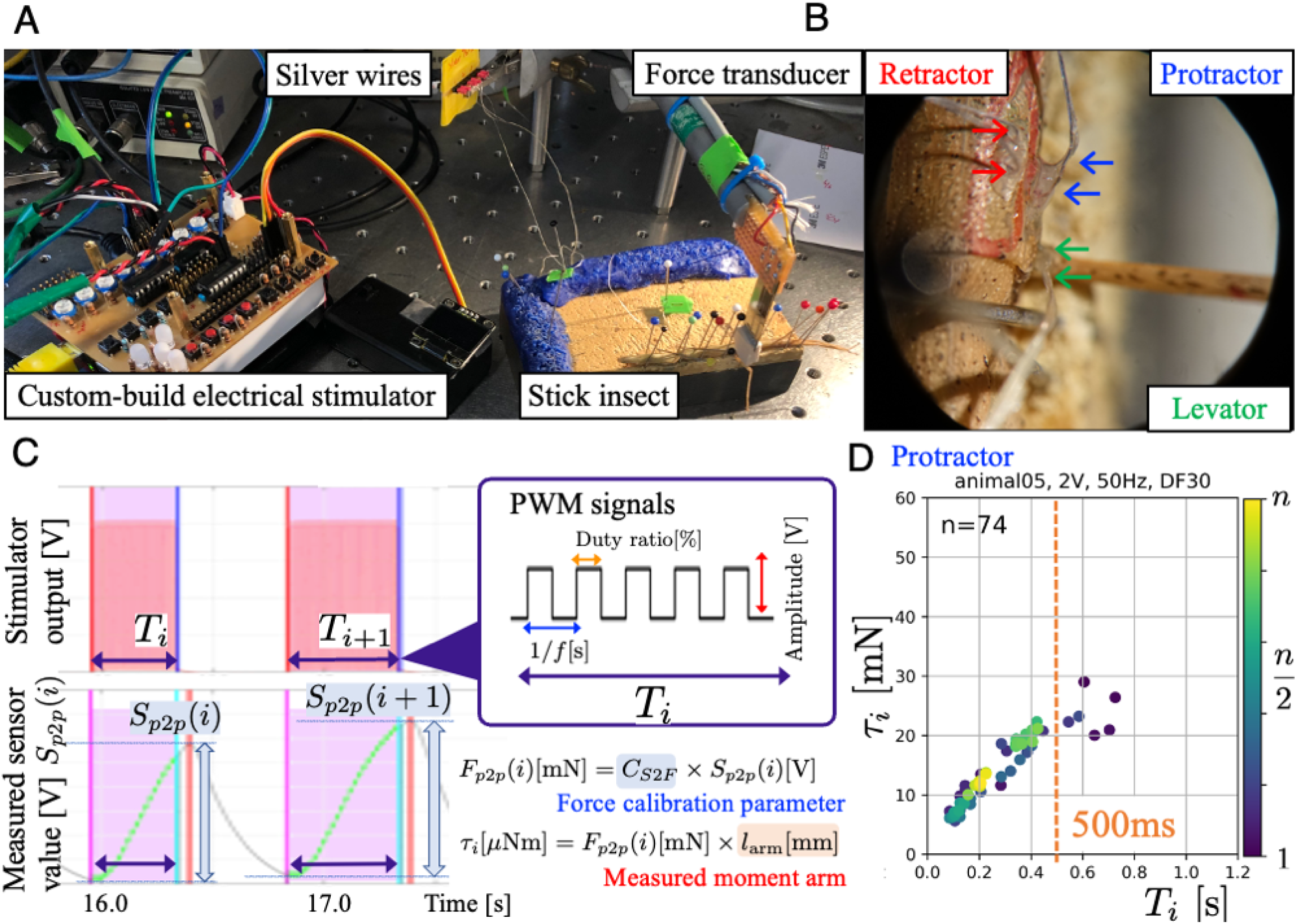
Experimental setup and joint torque calculation (**A**) The insect was fixed dorsal side up on a balsa wood platform. Two small insect pins attached to the tip of the force transducer held the middle part of the femur segment of the middle leg. (**B**) Electrodes (arrows) implanted into the three leg muscles protractor, retractor, and levator, in the right middle leg. (**C**) We systematically analyzed how joint torques depended on the three PWM burst parameters amplitude [V], frequency [Hz], and duty ratio [%], and identified the combinations that most effectively and repeatedly produced torque. The upper graph shows the profile of an electrical stimulation signal for each muscle. The lower graph shows the profile of the sensor value measured with the force transducer. (**D**). The panel shows the calculated ThC-joint torque profile versus the burst duration Ti during the protractor stimulations (animal05, *n* = 74). In this experiment, the burst duration Ti was varied at random, and the torque was calculated from force measurements with calibrated conversion factor and moment arm (see (**C**)). The voltage, frequency, and duty ratio of the PWM signals were 2.0 V, 50 Hz, and 30%, respectively. The color of the dots represents the number of stimulations (blue–yellow: 1–74). The orange dotted vertical line indicates *T_t_* at 500ms.

Finally, so far we have not fully investigated the effects of the electrical muscle stimulation on sensory feedback. The maximum voltage of 4V used here did not cause abnormal motion that could be attributed to cross-talk stimulation of sensory afferents. Therefore we conclude that unintended electrical stimulation of sensory afferents was negligible. Moreover, control measurements confirmed that muscles other than those stimulated by the electrodes were not active and did not generate force, as it would be expected from unintended stimulation via cross-talk. More generally, the activation of sensory organs during cyborg control is an interesting topic, with strong potential for expanding the concept of Motion Hacking. In the future, we will examine the performance of external leg movement control in an experimental setup, both without load (i.e., on a tether, without substrate contact) and with natural load distribution (i.e., by intervention during free walking). We are confident that these experiments, will provide further support of the Motion Hacking method and will reveal findings that could not be obtained by more conventional experiments without external stimulation of the neuro-muscular system. This will also contribute to potential applications in highly precise insect cyborg control.

## Methods and Materials

### Animals

We tested 20 adult female *Carausius morosus* from our laboratory colony at Bielefeld University in 2018. The animals were raised under a 12 h:12 h light:dark cycle at 23.9 ± 1.3 °C (mean ± S.D). Experiments were performed at a room temperature of 20–24 °C.

### Experimental setup

The insect was fixed dorsal side up on a balsa wood platform, using insect pins. The coxa of the right middle leg was located at the platform edge (Fig. 5 A right). We selected three leg muscles (protractor, retractor, and levator) in the right middle leg for electrical stimulation (Fig. 5 B). When stick insects walk, they use the protractor to swing the leg forward during the swing phase, the retractor to move the leg backward during the stance phase, and levator to initiate the stance-to-swing transition (***Rosenbaum et al., 2010***; ***Dallmann et al., 2019***; ***Günzel et al., 2022***; ***Bässler and Wegner, 1983***). Moreover, co-contraction of the protractor and retractor are known to vary based on the overall load distribution, thus being important for postural control by regulating joint stiffness (***Dallmann et al., 2019***; ***Günzel et al., 2022***). Electrical stimulation of the protractor and retractor muscles generate forward and backward leg movements at the thorax–coxa (ThC) joint, whereas stimulation of the levator muscle generates an upward leg movement at the coxa–trochanter (CTr) joint (***Dallmann et al., 2016***). To estimate the joint torque generated during the stimulation, we used a custom-made force transducer with strain gauges. Prior to the experiments, the measured force [mN] was calibrated from the force-sensor value [V] with weights of known mass (0.2–5 g). Two small insect pins attached to the tip of the force transducer held the middle part of the femur of the middle leg (Fig. 5 A right). The length between the ThC or CTr joints and the attachment point at the femur was measured and used as the moment arm for the calculation of torque.

### Electrical stimulation

We developed a custom-built electrical stimulator for stimulating muscles (Fig. 5 A left). An extension circuit board was designed for Raspberry Pi 3 B+ (Raspberry Pi Foundation), including isolated 8-channel PWM signal outputs. The parameters of the PWM signals, for example, frequency (1 to 120Hz) and duty ratio (0 to 100%), were changed using a Raspberry Pi microprocessor. The amplitude of the output voltage (0 to 9V) was changed using variable resistors on the circuit board, which enabled the investigation of the effects of these parameters on torque generation due to muscle stimulation. In this study, we systematically analyzed the joint torques generated by muscle contraction as induced by bursts of PWM pulses. To do so, we varied the amplitude [V], frequency [Hz] and duty ratio [%] of the PWM-signal, and identified the combinations that most effectively and repeatedly produced torque.

For one trial of the stimulation experiments, the frequency, duty ratio, and amplitude (voltage) of the PWM signals were not changed, but the burst duration *T_i_* of the signals was changed (Fig. 5 C top). Owing to the slow activation dynamics of an insect muscle, burst duration is one of two key parameters for controlling isometric muscle-contraction force because the muscle essentially acts as a second-order low-pass filter 30. The pulse frequency is the other key parameter, which can be held constant because burst duration alone is sufficient to effectively control joint torques in the range of 0–1.0 [s].

### Electrode implementation

A pair of stimulation electrodes was implanted into each muscle through two small holes in the cuticle. Holes were pierced using an insect pin, and wires were fixed with dental glue (Fig. 5 B). The stimulation electrodes were thin silver wires (A-M Systems, diameter = 127 μm, without insulation; 178 μm with Teflon insulation). The insulation at the end of the silver wire was removed, and the wires were implanted. The other end of the stimulation electrode was connected to the output of the electrical stimulator. The correctness of the electrode implantation was verified through triggered *resistance reflexes*, which are responses to imposed movements of the ThC and CTr joints for the corresponding muscles.

### Data Analysis

To investigate the dependence of externally induced joint torques by electrostimulating one out of the three leg muscles, we measured the force generated at the attachment point and multiplied it with the known moment arm as follows: (1) For different burst duration *T_i_* we estimated peak-to-peak sensor values *S_p2p_*(*i*) [V] (Fig. 5 C left). (2) Applying the conversion factor obtained from the previous calibration, we obtained peak-to-peak force change [N] in response to stimulation. (3) The force was then multiplied with the measured moment arm to obtain the joint torque *τ_l_* [Nm] (Fig. 5 C right).

### Bayesian Statistical Modeling

To investigate joint-torque properties generated by muscle stimulation while explicitly considering inter-individual variation of muscle physiology, we used a Bayesian statistical analysis and modeling framework. The probabilistic nature of Bayesian models makes them appropriate for modeling “uncertainty,” as introduced by inter-individual variation (***Gelman et al., 2013***). Bayesian analysis can be used to estimate a probabilistic distribution (model) that encodes an unknown observation target by using observed data and updating the distribution in the model. Furthermore, hierarchical model variants allow the inclusion of a hyperparameter, thus allowing for a parameter of choice to be drawn from yet another probabilistic distribution. In our case, hierarchical-model variants were used to account for inter-individual differences (***Watanabe, 2018***).

Here, we modeled the relationship between the burst duration of the electrical stimulation and the joint torque generated using a single model (a power function) with six variants (for details, see subsection “Models”). All model variants were specified in a probabilistic programming language developed by Stan (***Stan Development Team, 2022***). Here, we used non-informative uniform priors for the parameters *β, γ*, and *σ*, unless stated otherwise. For estimation, we used the numerical Markov Chain Monte Carlo (MCMC) method, and scripted the models in R (v.4.1.1) (***R Core Team, 2022***), in which the Stan code was compiled and executed using the R package “rstan” (***Stan Development Team, 2022***). The software performed sampling from prior distributions using No-U-Turn Sampler (NUTS) (***Hoffman and Gelman, 2014***). Sampling convergence was detected through trace plots and the quantitative Gelman-Rubin convergence statistic *R_hat_*(***Gelman and Rubin, 1992***), where *R_hat_*< 1.10.

### Models

*τ_i_* and *T_i_* represent the calculated joint torque based on the force-transducer value and the burst duration of a PWM signal for electrical stimulation, respectively. We assumed that *τ_i_* follows a normal distribution, described by the 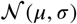 function, where *μ* and *σ* represent the mean and standard deviation (S.D.) of the distribution. Indexes *i* and *j* represent the numbers of stimulations and animals, respectively.

#### Model 1-1

Linear model representing the linear relationship between burst duration and joint torque

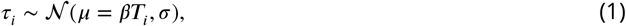

where *β* represents the inclination of the estimated linear function.

#### Model 1-2

Hierarchical model representing the linear relation between burst duration and joint torque

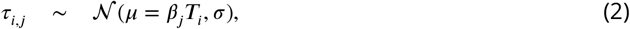

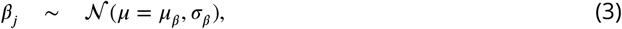

where *β_j_* represents the inclination of the estimated linear function on the (*j*)th animal. Furthermore, in this hierarchical model, *β_ĵ_* is drawn out of a normal distribution that captures inter-individual variation, where *μ_β_* and *σ_β_* represent the mean and S.D. of the distribution, respectively.

#### Model 2-1

Non-linear model representing the nonlinear relationship between burst duration and joint torque

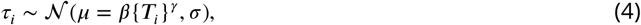

where, *β* and *γ* represent the magnitude of the base and exponent of the estimated non-linear power function, respectively.

#### Model 2-2

Hierarchical model representing the nonlinear relationship between burst duration and joint torque

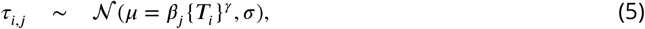

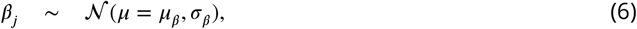

where *β_j_* and *γ* represent the magnitude of the base on the (*j*)th animal and the exponent of the estimated nonlinear power function, respectively. In this model, *β_j_* follows a normal distribution as described above, where, *μ_β_* and *σ_β_* represent the mean and S.D. of the distribution.

#### Model 2-3

Hierarchical model representing the nonlinear relation between burst duration and joint torque

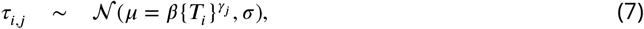

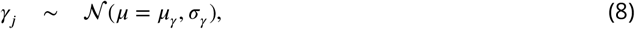

where *β* and *γ_j_* rrepresent the magnitude of the base and the exponent on the (*j*)th animal for the estimated nonlinear, power function, respectively. In this model, *γ_β_* follows a normal distribution as described above, where *μ_γ_* and *σ_γ_* represent the mean and S.D. of the distribution.

#### Model 2-4

Hierarchical model o representing the nonlinear relationship between burst duration and joint torque

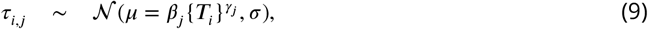

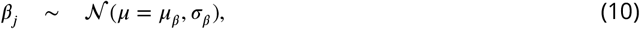

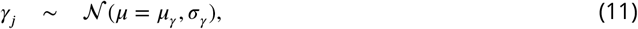

where *β_j_* and *γ_j_* represent the magnitude of the base and the exponent of the estimated nonlinear power function, respectively, on the (*j*)th animal. In this model, *β_j_* and *γ_j_* follow normal distributions as described above, where *μ_β_* and *μ_β_* are the means, and *σ_γ_* and *σ_β_* are the S.D.s of the distribution.

### Widely Applied Information Criterion (WAIC)

We compared the predictive performances of the formulated models using the mathematical index WAIC (***Watanabe, 2018, 2005, 2010a,b***). The WAIC is a measure of the degree to which an estimate of the predictive distribution is accurate relative to the true distribution (***Watanabe, 2018***). Essentially, it is based on the difference between the information conveyed by the mean and that conveyed by the variance. This difference is negative if the term corresponding to the mean exceeds that corresponding to the variance. The smaller (or more negative) the WAIC index, the higher the predictive value of the model variant.

When calculating the WAIC index for a hierarchical model, several calculation methods can be used, depending on the definition of the predictive distribution, that is, the type of unknown data distribution being predicted (***Watanabe, 2018***). We were interested in predicting muscle properties with electrostimulation for a new, additional animal, not including experimental date, to enable individualized leg “control.” From this perspective, we constructed a new distribution of the predictive parameters of a new animal by marginalizing intermediate parameters assigned to each hierarchical model (models 1-2, 2-2, 2-3, 2-4) (***Watanabe, 2018***; ***Wakita et al., 2020***; ***Harada et al., 2020***). This allows for a fair comparison of the prediction performance of hierarchical and non-hierarchical models. Referring to the method from the previous studies (***Wakita et al., 2020***; ***Harada et al., 2020***), the WAIC was computed by numerical integration with MCMC (Markov Chain Monte Carlo) samples by using Simpson’s law and the “log_sum_exp” function provided by Stan (***Stan Development Team, 2022***).

From the models described above, the model with the smallest WAIC value was considered the most appropriate predictive model in terms of predictivity for a new animal.

## Acknowledgments

This work was supported by JSPS KAKENHI Grant-in-Aid for Scientific Research on Innovative Areas “Science of Soft Robot” project under Grant Number JP21H00317 and a Grant-in-Aid for the Promotion of Joint International Research (Fostering Joint International Research) (JP17KK0109).

## Data Availability

We will share a database of all experimental data in the final version of the paper on Dryad (https://datadryad.org/). Experimental device information (circuit design, etc.) and codes also will be available on repositories such as GitHub.

## Ethics statement

At present, animal care regulations does not need to be considered in our insect research in Bielefeld University and Tohoku University. However, the authors totally agreed that future research on cyborg insects, which pushes boundaries not yet fully considered by ethicists and legislators, will require careful ethical consideration of both animal welfare and social consequences.

## Author contributions

Dai Owaki, Conceptualization, Data curation, Formal analysis, Funding acquisition, Investigation, Methodology, Project administration, Resources, Software, Validation, Visualization, Writing – original draft, Writing – review and editing; Volker Dürr, Conceptualization, Investigation, Resources, Supervision, Validation, Writing – review and editing; Josef Schmitz, Methodology, Data curation, Investigation, Resources, Supervision, Project administration, Validation, Writing – review and editing

